# Resegmentation is an ancestral feature of the gnathostome vertebral skeleton

**DOI:** 10.1101/779710

**Authors:** Katharine E. Criswell, J. Andrew Gillis

## Abstract

The vertebral skeleton is a defining feature of vertebrate animals. However, the mode of vertebral segmentation varies considerably between major lineages. In tetrapods, adjacent somite halves recombine to form a single vertebra through the process of “resegmentation”. However, in teleost fishes, there is considerable mixing between cells of the anterior and posterior somite halves, without clear resegmentation. To determine whether resegmentation is a tetrapod novelty, or an ancestral feature of jawed vertebrates, we tested the relationship between somites and vertebrae in a cartilaginous fish, the skate (*Leucoraja erinacea*). Using cell lineage tracing, we show that skate trunk vertebrae arise through tetrapod-like resegmentation, with anterior and posterior halves of each vertebra deriving from adjacent somites. We further show that tail vertebrae also arise through resegmentation, despite a duplication of the number of vertebrae per body segment. These findings resolve axial resegmentation as an ancestral feature of the jawed vertebrate body plan.

## Introduction

Axial segmentation is key to the body plan organization of many metazoan groups and has arisen repeatedly throughout animal evolution (Davis and Patel, 1999). Within vertebrates, the axial skeleton is segmented into repeating vertebral units that provide structural support and protection for soft tissues. Vertebral segmentation is preceded in the embryo by the segmentation of paraxial mesoderm into epithelial blocks called somites (Figure 1a). Cells from the ventromedial portion of each somite then undergo an epithelial to mesenchymal transition and migrate around the notochord and neural tube, where they condense into vertebrae. Cell lineage tracing studies in tetrapods have shown that there is not a 1:1 correspondence between somites and vertebrae. Rather, cells from adjacent somite halves recombine to give rise to a single vertebra, through a process known as “resegmentation” (Remak, 1855; Verbout, 1976). Somite lineage tracing in chick using chick-quail chimeras or lipophilic dyes (Aoyama and Asamoto, 2000; Bagnall et al., 1988; Huang et al., 2000; Schrägle et al., 2004; Stern and Keynes, 1987; Ward et al., 2017) has shown that cells from the rostral half of one somite combine with cells from the caudal half of the adjacent somite to give rise to a single vertebra, with sharp compartment boundaries in the middle of vertebrae reflecting original somite boundaries. Additionally, lineage tracing experiments in axolotl using injections of fluorescent dextrans or grafts of GFP+ somites into wildtype hosts point to conservation of resegmentation during vertebral development in lissamphibians (Piekarski and Olsson, 2014). However, in teleost fishes, resegmentation is less apparent, with cells from adjacent somite halves undergoing substantial mixing, resulting in vertebrae without clear lineage-restricted compartments (Morin-Kensicki et al., 2002). This divergence in the relationship between somites and vertebrae raises the question of whether resegmentation is restricted to tetrapods, or whether it is an ancestral feature of the jawed vertebrate (gnathostome) backbone that has been reduced/lost in teleosts. To answer this question, data are needed from an outgroup to the bony fishes, the cartilaginous fishes.

**Figure 1.**
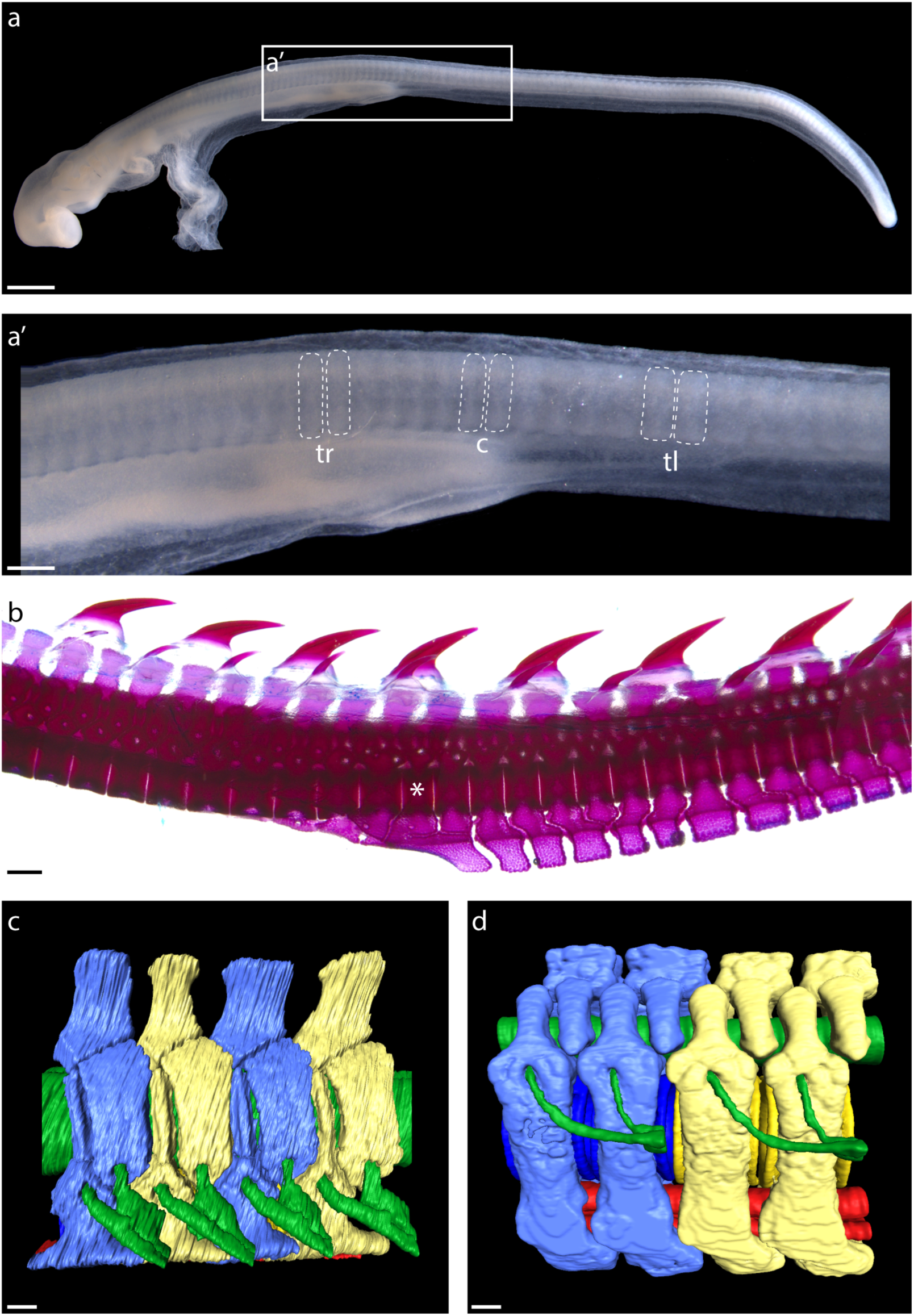
Segmental organization of paraxial mesoderm and the vertebral skeleton in the skate. **a)** Somites of a S24 skate embryo showing **a’)** trunk (tr), cloacal (c), and tail (tl) somites, delineated with dashed lines. **b)** A cleared and stained vertebral column from a skate hatchling, with the trunk to tail transition denoted with an asterisk. **c)** µCT scan reconstruction of skate trunk and **d)** tail vertebrae, with vertebral elements corresponding to a single body segment color-coded blue or yellow, neural tissue colored green, and blood vessels colored red. Note that in **d)**, one set of spinal nerves spans two vertebrae. Scale bars: a) 400 µm; b) 100 µml c) 500 µm; d) and e) 100 µm.

The vertebral skeleton in cartilaginous fishes consists of a series of neural and intercalary arches and neural spines, tri-layered centra and haemal arches (the latter restricted to the caudal region – Figure 1b) (Criswell et al., 2017a). Notably, the axial skeleton of the embryonic skate forms initially as a continuous, sclerotome-derived cartilaginous tube, which subsequently subdivides into discrete vertebrae (Criswell et al., 2017a, 2017b). Elasmobranch cartilaginous fishes (sharks, skates and rays) also show a unique pattern of spinal nerve segmentation along the body axis, with each trunk vertebra containing foramina for both dorsal and ventral spinal nerve roots, while each tail vertebra contains a foramen for only a single nerve root (Ridewood, 1899) (Figure 1c,d). This pattern is widely hypothesized to reflect a complete vertebral duplication in each tail body segment, termed “diplospondyly”. However, there are currently no experimental data testing this, or, more generally, the relationship between somites and vertebrae, in any cartilaginous fish.

Here we test whether tetrapod-like resegmentation is the ancestral mode of vertebral segmentation in gnathostomes using an outgroup to the bony fishes, a cartilaginous fish (the little skate, *Leucoraja erinacea*). We also test whether the diplospondyly observed in the tails of cartilaginous fishes arises through duplication of skeletal derivatives from tail vs. trunk somites. We find that skate tail somites do, indeed, duplicate their vertebral contribution relative to trunk somites, and that resegmentation occurs along the entire body axis. These findings point to a stem-gnathostome origin of axial column resegmentation, and allow us to reconstruct the evolutionary history of vertebral column development.

## Results

### Conservation of rostrocaudal somite polarity in tetrapods and skate

Tetrapod somites exhibit a distinct rostrocaudal polarity, for example, with expression of the T-box transcription factor *Tbx18* localizing to the rostral somite halves, and expression of the homeobox transcription factor *Uncx4*.1 localizing to the caudal somite halves (Haenig and Kispert, 2004; Kraus et al., 2001; Schrägle et al., 2004). These distinct rostral and caudal molecular identities are essential for proper resegmentation (Bussen et al., 2004; Hughes et al., 2009; Keynes, 2018; Leitges et al., 2000; Mansouri et al., 2000). To test for rostrocaudal polarity of somites in skate, we examined expression patterns of *Tbx18* and *Uncx4.1* using wholemount mRNA *in situ* hybridization. We found that *Tbx18* (Figure 2a) and *Uncx4.1* (Figure 2b) are both expressed in skate somites at S22, with expression of the former localizing to the rostral somite, and expression of the latter localizing to the caudal somite. To further test polarity of these expression patterns within the somite, we characterized transcript distribution on paraffin sections of somites using multiplexed fluorescent mRNA *in situ* hybridization, and we found that *Tbx18* and *Uncx4.1* transcripts localize to the rostromedial and caudomedial cells of the somite, respectively (Figure 2c). These findings point to a shared molecular basis of somite rostrocaudal polarity between tetrapods and cartilaginous fishes.

**Figure 2.**
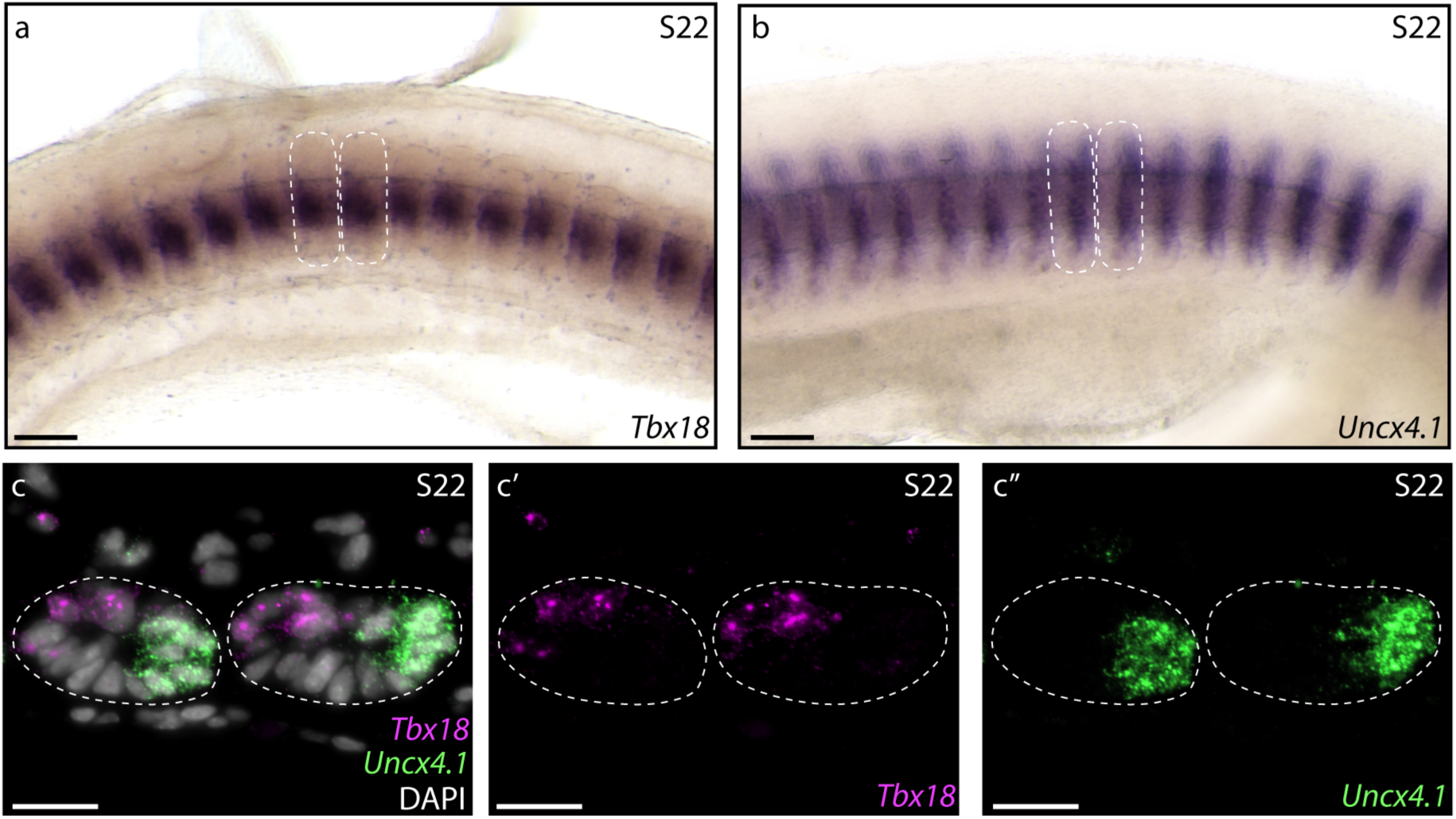
Molecular basis of skate somite rostrocaudal polarity. Whole-mount mRNA *in situ* hybridization reveals **a)** rostral expression of *Tbx18* and **b)** caudal expression of *Uncx4.1* within skate somites at S22. **c)** Hybridization chain reaction (HCR) *in situ* hybridization on paraffin sections reveals complimentary rostromedial and caudomedial expression of *Tbx18* (magenta, **c’**) and *Uncx4.1* (green, **c’**’), respectively, in skate somites at S22. Dashed lines indicate somite boundaries. Scale bars: a) and b) 100 µm; c-c’’) 25 µm.

### Skates exhibit tetrapod-like resegmentation

To test the relationship between somites and vertebrae in skate, we performed a series of somite fate mapping experiments. We microinjected the ventral portions of two neighboring trunk somites in S24 little skate embryos with either CM-DiI (a red fluorescent analogue of DiI) or SpDiOC_18_ (a green fluorescent analogue of DiO) (Figure 3a), and then mapped the contributions of these labeled somites to the vertebral skeleton 8-12 weeks post-injection. We recovered dye in the vertebral cartilage of 27 embryos, with a clear offset between somite and vertebral boundaries in all instances. 21/27 embryos retained both CM-DiI and SpDiOC_18_ labelling in adjacent vertebrae, while 6/27 retained only CM-DiI. In the trunk, two neighboring somites contributed to a combined total of three vertebrae, with each somite contributing to two vertebral halves (the caudal half of one vertebra and the rostral half of the neighboring vertebra - Figure 3b-d; n=22/27). There was little overlap in CM-DiI and SpDiOC_18_ distributions, indicating that cells from adjacent somites undergo little mixing, and that somite boundaries are maintained through sclerotome migration, and condensation. The results of these labeling experiments are consistent with a tetrapod-like mechanism of resegmentation during the development of trunk vertebrae in the skate.

**Figure 3.**
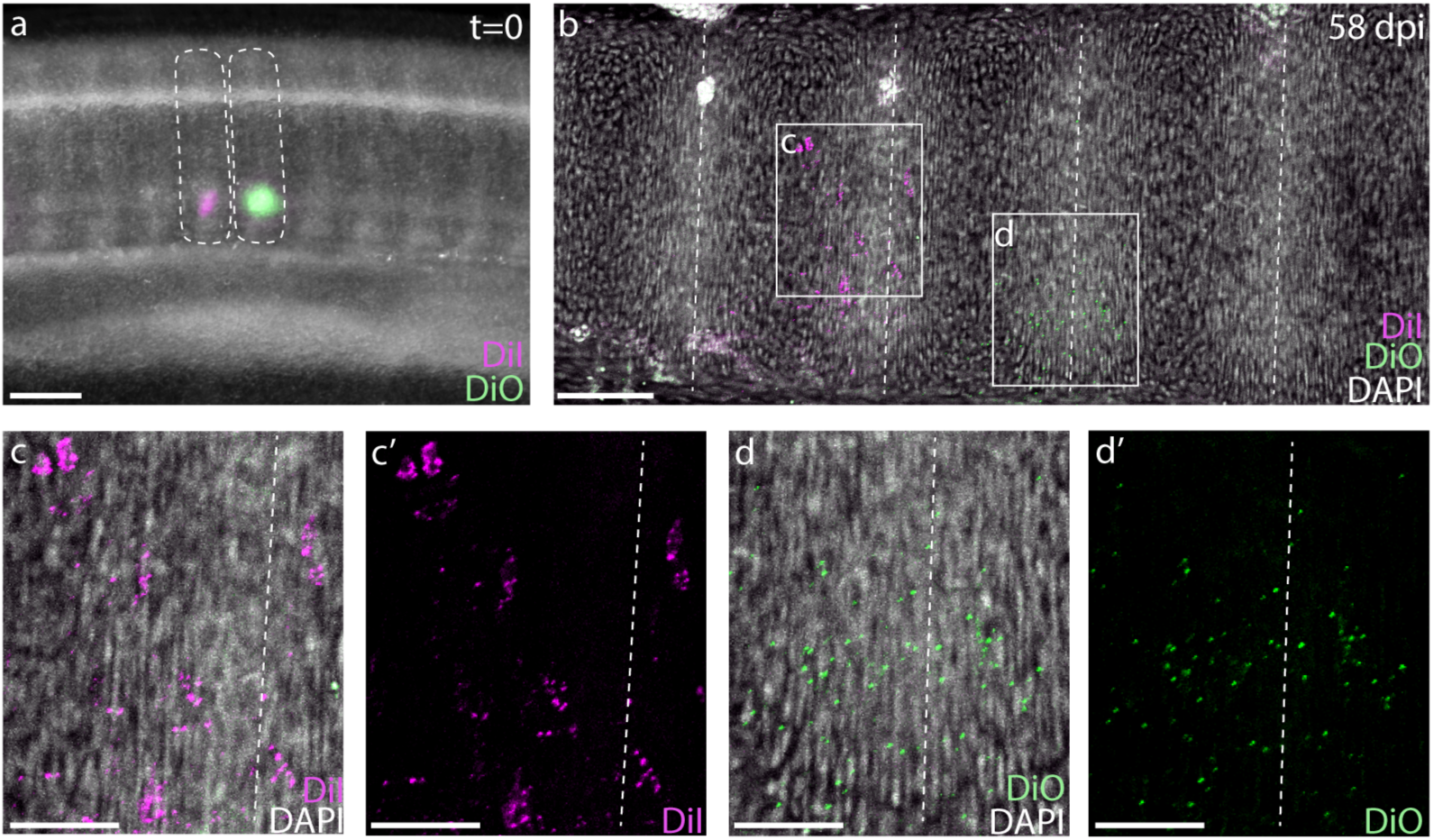
Skates undergo tetrapod-like resegmentation. **a)** Adjacent CM-DiI- and SpDiOC_18_-labeled trunk somites in a S24 skate embryo. **b)** Sagittal section of the trunk of an embryo 58 days post-injection, showing CM-DiI- and SpDiOC_18_-labeled cells in the posterior and anterior halves of adjacent vertebrae, consistent with resegmentation. **(c, c’)** CM-DiI-labeled cells and **(d, d’)** SpDiOC_18_-labeled cells span two vertebrae (each shown with and without DAPI counterstain). Dashed lines indicate somite boundaries in **(a)** and vertebral boundaries in (**b-d)**. Scale bars for **a)** and **b)** 100 µm; scale bars for **c)**-**d’)** 50 µm.

### Skate tail somites give rise to twice as many vertebrae as trunk somites, and still undergo resegmentation

Elasmobranch fishes show a diplospondylous condition in their caudal vertebrae, in which two vertebrae are present for each set of spinal nerve roots, and it is speculated that this condition is a consequence of tail vertebrae giving rise to twice as many vertebral units when compared with trunk vertebrae. If somites give rise to double the number of vertebrae in the tail as in the trunk, we would expect approximately equal numbers of trunk somites and vertebrae, while the number of tail vertebrae should far exceed the number of tail somites. We counted trunk and tail somites in S25 skate embryos (at which point somitogenesis has ceased, but somites are still clearly discernable), and compared this with numbers of trunk and tail vertebrae in skate hatchlings. We found that S25 skate embryos possess a mean of 48 trunk somites and 88 tail somites (with the cloaca marking the transition from trunk to tail; n=17 embryos counted; Supplementary Table 1), while hatchling skates possess a mean of 48 trunk vertebrae (including the series of fused vertebrae that make up the synarcual, which was determined by counting sets of spinal nerve foramina) and 104 tail vertebrae (n=8 hatchlings counted; Supplementary Table 2). These counts are consistent with the hypothesis that tail somites give rise to more (though not consistently double the number of) vertebral units compared with trunk somites. It should be noted, however, that there is difficulty in accurately counting terminal vertebrae in skates: the tail tapers to a fine point, with terminal vertebrae becoming extremely small and difficult to differentiate in skeletal preparations, and it is also likely that additional nons-mineralized vertebral elements are present in the tip of the tail that we were not able to distinguish by this method. We therefore speculate that a pattern of duplication of somite derivatives observed in the anterior tail may not persist along the full length of the tail, but rather breaks down near the tip.

To test whether a single somite gives rise to more than two vertebral halves in the presumptive diplospondylous vertebrae of the skate tail – and to assess whether the process of somite segmentation occurs in the tail as in the trunk – we repeated the above somite labeling experiments in adjacent somites posterior to the cloaca (Figure 4a). Analysis of labeled embryos 10 weeks post-injection revealed CM-DiI and SpDiOC_18_ each in the cartilage of three vertebrae: the posterior portion of one, the entire next successive vertebra, and the anterior portion of the third (n=19/23, Figure 3f-h). Each tail somite therefore contributes to parts of five vertebrae, fully double that of each trunk somite, and again consistent with a tetrapod-like model of somite resegmentation.

**Figure 4.**
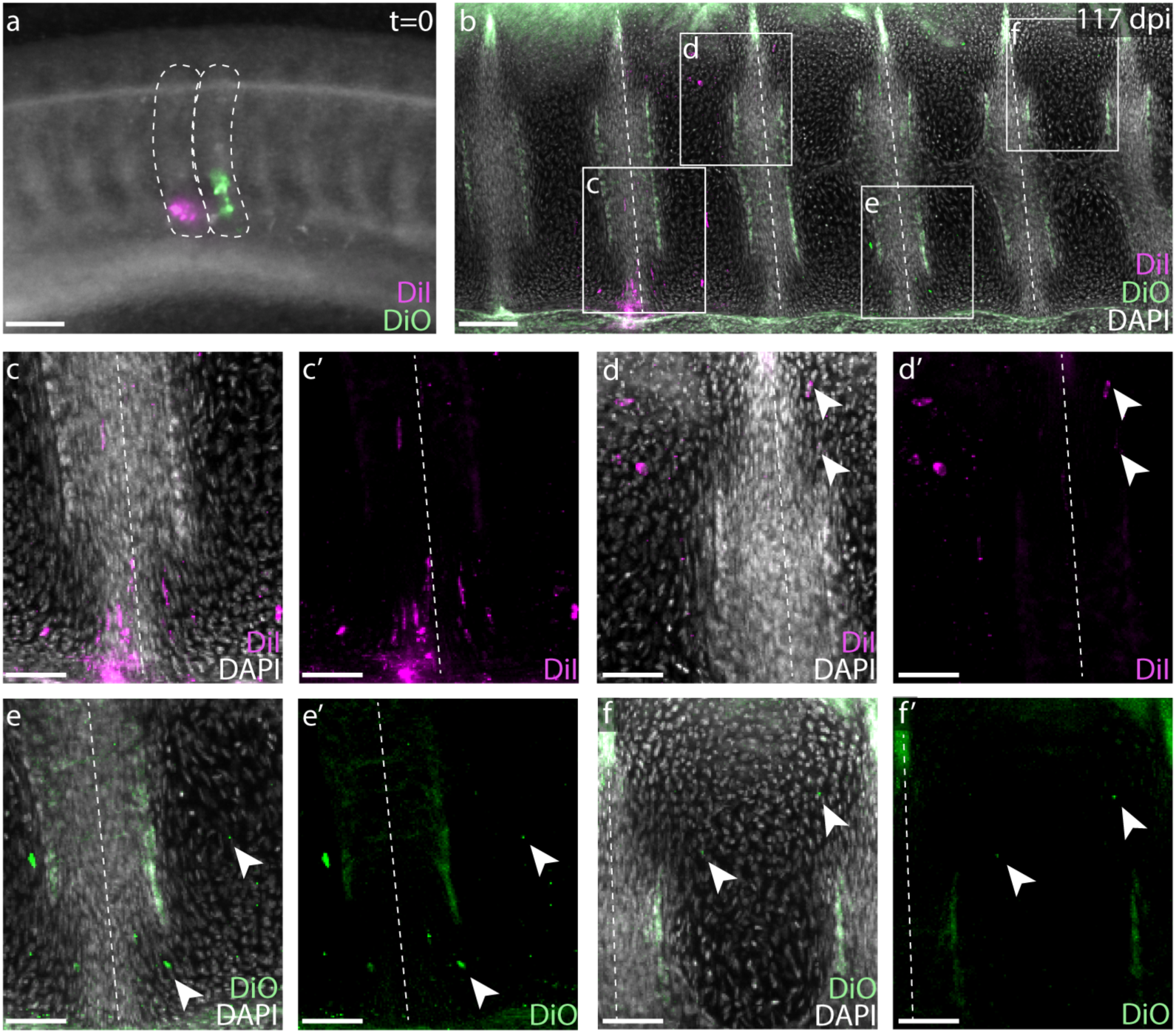
Skate tail vertebrae are duplicated and resegmented. **a)** Adjacent CM-DiI and SpDiOC_18_ injections in tail somites of a S24 little skate embryo. **b)** Sagittal section of the tail of an embryo 117 days post-injection showing CM-DiI- and SpDiOC_18_-labeled cells in three vertebrae each. Magnified views reveal **c)-d)** CM-DiI- and **e)-f)** SpDiOC_18_-labeled cells spanning vertebral boundaries (images shown with without DAPI counterstain). Dashed lines indicate somite boundaries in **(a)** and vertebral boundaries in **(b-f)**. Arrowheads indicate dye-labeled cells. Scale bars for **a)** and **b)** 100 µm; scale bars for **c)**-**f’)** 50 µm.

Because the reduction in spinal nerve foramina per vertebral segment coincides with the first caudal vertebra (Figure 1b), we hypothesized that the transition between the monospondylous vertebrae of the trunk and the diplospondylous vertebrae of the tail occurred at the cloaca. To test the location of this transition, we injected one of the posterior-most somites just dorsal to the cloacal opening with CM-DiI (Figure 5a). At 8-12 weeks post-injection, most cloacal-region somite injections resulted in the trunk-like resegmentation pattern of two vertebral halves derived from one somite (Figure 5b;c, n=8/9). In each case, CM-DiI-labeled chondrocytes were recovered in vertebrae within several segments of the monospondylous-diplospondylous transition, suggesting that this boundary occurs just posterior to the cloaca.

**Figure 5.**
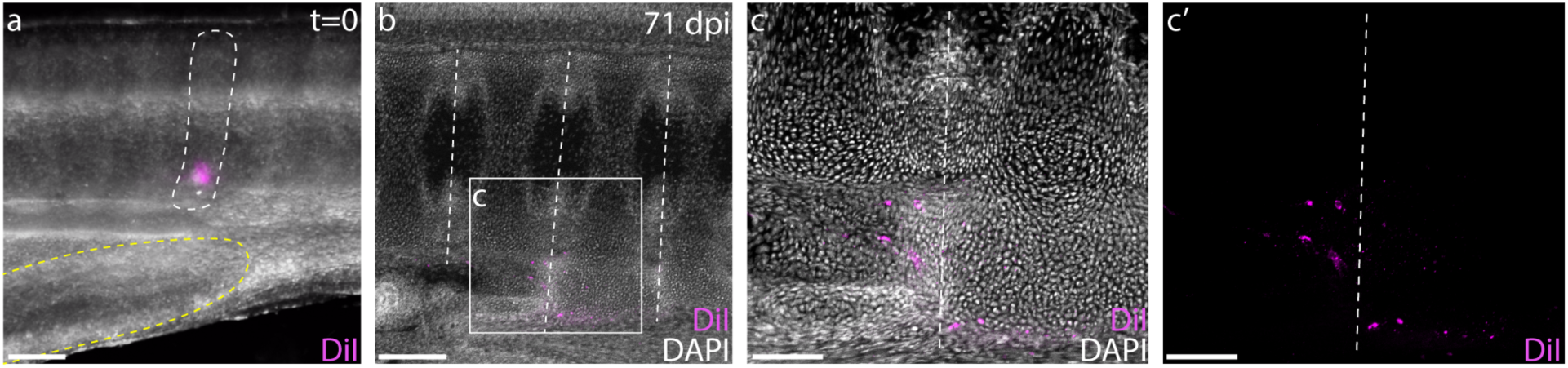
The cloaca precedes the monospondylous to diplospondylous transition. **a)** CM-DiI labeling of a S24 skate somite (marked with white dashed line) just dorsal to the posterior margin of the cloaca (denoted with yellow dashed line). **b)** Sagittal section of an embryo 71 days post-injection, showing two adjacent vertebrae labeled with CM-DiI. **c)** Magnified view of two vertebrae showing CM-DiI labeled cells with and **c’)** without DAPI counterstain. Dashed lines in **b)**-**c)** indicate vertebral boundaries. Scale bars for **a)** and **c)** 100 µm; scale bars for **b)** 200 µm.

## Discussion

Here we report the presence of strict somite resegmentation in a cartilaginous fish. Our two-color fate mapping experiments show clearly that a single skate trunk somite gives rise to two adjacent vertebral halves. When combined with fate mapping data from axolotl and chick, our findings indicate that vertebrae were formed through somite resegmentation in the last common ancestor of jawed vertebrates, and that the intermixing of somite cells throughout vertebral bodies seen in zebrafish represents a departure from that ancestral pattern (Figure 6a). We also show that resegmentation is maintained along the AP axis in skate, despite differences in the number of somite derivatives in the trunk and tail. In the trunk, each somite gives rise to two vertebral halves spanning a single vertebral boundary (Figure 6b), and this transitions posterior to the cloaca (Figure 6c) to tail somites giving rise to four vertebral halves spanning two vertebral boundaries (i.e. a single tail somite gives rise to a half vertebra, a full vertebra, and a subsequent half vertebra – Figure 6d). Unlike the ancestral pattern of resegmentation in the skate trunk, the modification of this arrangement to give rise to duplicated vertebrae in the tail is unique to elasmobranchs and, to our knowledge, previously unreported.

**Figure 6.**
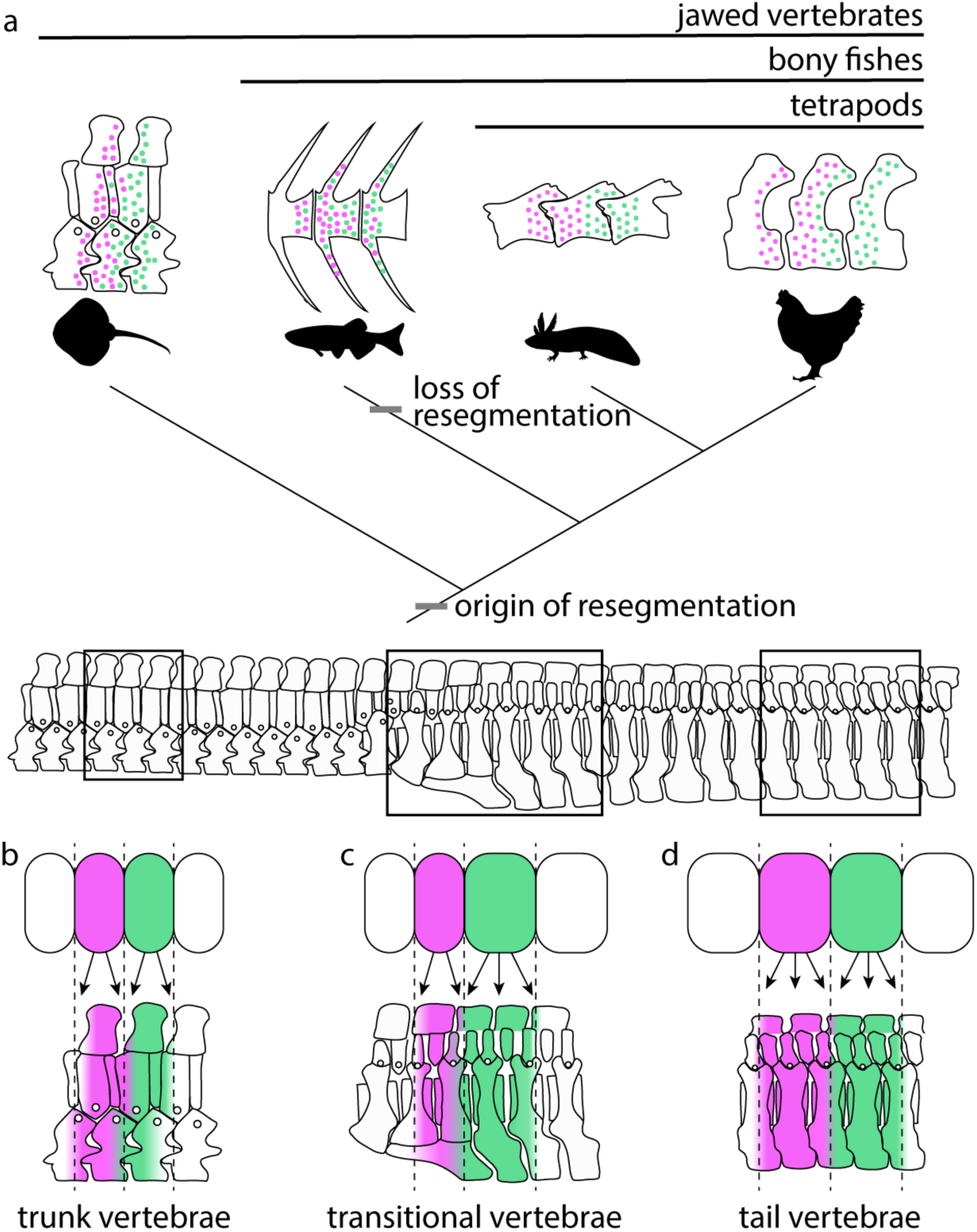
Resegmentation is ancestral for jawed vertebrates. a) Schematic showing the distribution of strict resegmentation on a simplified vertebrate evolutionary tree, and the contribution of two adjacent somites to b) trunk vertebrae; c) vertebrae at the trunk-to-tail transition; and d) tail vertebrae in the skate.

Our demonstration of a shared molecular basis of rostrocaudal somite polarity between tetrapods (Bussen et al., 2004; Haenig and Kispert, 2004; Kraus et al., 2001; Schrägle et al., 2004), teleosts (Begemann et al., 2002), and skate suggests that this feature is likely also ancestral for jawed vertebrates, and probably a precondition for resegmentation. Quail-chick chimera experiments have shown that only sclerotome derived from like somite halves will intermingle following recombination – e.g. donor cells derived from rostral half-somites will intermingle with neighboring host sclerotome only when grafted next to another rostral half-somite, but will obey a tight boundary with no cell mixing when grafted next to a caudal half-somite (Stern and Keynes, 1987). This tendency for like cells to form discrete compartments within the somite likely underlies the subsequent contribution of rostral somite cells to caudal vertebrae, and vice versa, upon resegmentation. Interestingly, if this strict compartmentalization of somite halves is shared with skate, our finding of resegmentation in the diplospondylous tail region would suggest that rostral and caudal somite halves are not committed to caudal and rostral vertebral fates, respectively, but rather have the capacity to differentiate into either (as we would predict that the rostral half of a single somite would contribute to the caudal half of one vertebra and the rostral half of a second, neighboring vertebra, while the caudal half-somite would contribute to the caudal half of the second vertebra and the rostral half of a third). However, more specific lineage tracing experiments mapping the fates of half-somites are needed to test this.

*Tbx15/18/22* and *Uncx4.1* are expressed in the rostral and caudal halves, respectively, of some somites in the invertebrate chordate amphioxus, in a pattern similar pattern to that of vertebrates (Beaster-Jones et al., 2008). This indicates that the molecular basis of somite AP polarity likely arose along the chordate stem, preceding the evolutionary origin of vertebrae. *Tbx15/18* is expressed in anterior somite halves in lamprey (Freitas et al., 2006), which also possess a vertebral skeleton in the form of small, rudimentary neural arches. Lineage tracing experiments in lamprey to track the contribution of single somites to the neural arch cartilages would help to further resolve the origin of resegmentation within vertebrates.

Historically, resegmentation was hypothesized to have originated through a functional need for the staggered positioning of myotomes and vertebrae in aquatic vertebrates. The articulation of muscle fibers from one myotome across a vertebral joint to two individual vertebral centra was thought to facilitate lateral bending and therefore axial locomotion (Lauder, 1980). However, the relative positioning of arches to centra, and of centra within myomeres, can vary substantially between different actinopterygian species, and within species, along the anteroposterior axis (Schaeffer, 1967). Myosepta in jawed vertebrates also are morphologically highly complex, with a W-shape and six tendons that attach laterally to the skin and medially to the vertebrae and median septum, providing connections across up to three individual vertebrae (Gemballa S. et al., 2003; Gemballa and Röder, 2004). These anatomical features combined suggest that a positional frameshift resulting from strict resegmentation is not necessary for axial locomotion in non-amniote vertebrates. Our results, in which caudal somites undergo resegmentation, but not in register with individual vertebrae, support the conclusion that resegmentation is not necessary for axial locomotion. Whether any actual locomotory advantage was gained from the evolution of resegmentation remains unclear.

## Materials and Methods

### Animal collection and husbandry

All skate embryos were obtained from captive brood stock at the Marine Biological Laboratory in Woods Hole, Massachusetts, USA and all experimental work was conducted in accordance with approved IACUC protocols. Embryos were reared in flow-through seawater tables at 10-12°C for approximately four weeks prior to experimentation at stage (S) 24. Early skate embryos were staged according to Ballard et al. (1993) and late-stage embryos were staged following Maxwell et al. (2008).

### µCT scanning

Portions of the trunk and tail of a S34 skate embryo were stained with iodine potassium iodide (IKI) according to Metscher (2009), and scanned using a GE v|tome|x µCT scanner at the University of Chicago. The trunk was scanned at 80 kV and 70 uA, with an exposure time of four seconds and a voxel size of 3.763 µm. The tail was scanned at 100 kV and 100 uA with a two second exposure and a voxel size of 3.075 µm. CT slices were processed and segmented in Avizo (ThermoFisher Scientific - FEI). Tiff stacks for each scan are available on the Dryad digital repository, doi:10.5061/dryad.b2rbnzs8s.

### Somite and Vertebral counts

Vertebrae in hatchling skates were visualized by skeletal preparation, as described in Gillis et al. (2009) with an added overnight incubation in Alizarin red solution (1mg/mL Alizarin red in 1% KOH) and overnight trypsin (1% w/v in water) digestion prior to KOH clearing. Somites in S25 skate embryos (n=17) and vertebrae in cleared and stained skate hatchlings (n=8) were imaged in numerous focal planes on a Leica M165 FC stereoscope. Image stacks were then merged in Helicon Focus Pro and tiled to form high resolution images. Somites and vertebrae were counted in Adobe Photoshop 2018 using a layered overlay and dots to mark individual elements.

### Histology and mRNA in situ hybridization

*L. erinacea* embryos were embedded in paraffin wax and sectioned at 8 µm thickness for mRNA *in situ* hybridization as described in O’Neill et al (2007). Chromogenic mRNA *in situ* hybridization experiments for *Uncx4.1* (GenBank accession number MN478366) and *Tbx18* (GenBank accession number MN478367) were performed on sections as described in O’Neill et al. (2007), with modifications according to Gillis et al. (2012). Probes, buffers, and hairpins for third generation *in situ* hybridization chain reaction (HCR) experiments were purchased from Molecular Instruments (Los Angeles, California, USA). Experiments were performed on paraffin sections according to the protocol of Choi et al. (2018), with the following modifications: Following proteinase K treatment and rinsing, slides were pre-hybridized for 30 minutes at 37C, and then hybridized overnight at 37C with 0.8uL of 1uM probe stock/100uL of hybridization solution. Following post-hybridization washes and pre-amplification steps, slides were incubated in amplification solution containing 4uL of each hairpin stock/100uL of amplification buffer.

### Fate mapping experiments

Two-color somite fate mapping experiments were performed as described in Criswell et al (2017b) and Ward et al (2017). S24 skate embryos were removed from their egg cases to a petri dish and anesthetized in tricaine (MS-222 1 mg/L in seawater). Adjacent somites were injected with the red-fluorescent lipophilic dye CM-DiI and the green-fluorescent lipophilic dye SpDiOC_18_ (ThermoFisher). Concentrated stocks of CM-DiI (5 µg/µL in absolute ethanol) and SpDiOC_18_ (2.23 µg/µL in dimethylformamide) were diluted 1:10 in 0.3 molar sucrose for injection. After injection embryos were returned to their egg cases and maintained in a flow-through seawater table at 15°C for 8-12 weeks post-injection.

Injected skate embryos were euthanized using an overdose of tricaine (1g/L in seawater) and fixed in 4% paraformaldehyde overnight at 4°C. Embryos were then rinsed 3×5 minutes in phosphate buffered saline (PBS), embedded in 15% gelatin in PBS and post-fixed in 4% paraformaldehyde in PBS for 4 nights at 4°C before vibratome sectioning. A Leica VT1000S vibratome was used to cut 100 µm sections of tissue in sagittal plane, which were then DAPI-stained (1µg/mL), coverslipped with Fluoromount-G (Southern Biotech) and imaged on an Olympus FV3000 (trunk and transitional vertebrae) or Leica Sp5 (tail vertebrae) confocal microscope.

## Acknowledgements

We thank Dr Victoria Sleight, Christine Hirschberger, and Jenaid Rees, along with the Cambridge Evolution and Development community for valuable feedback. We are grateful to Dr. Richard Schneider, Prof. David Sherwood, and the MBL Embryology Course for provision of lab space, Louise Bertrand and Leica Microsystems for microscopy support, and the staff of the Marine Resources Center at the MBL for aid in animal husbandry. This project benefited from use of the Imaging Facility, Department of Zoology, supported by a Sir Isaac Newton Trust Research Grant (Ref 18.07ii(c))

## Supplemental Information

**Table S1.** Counts of trunk and tail somites in S25 skate embryos.

|  | **Trunk**<br>**somites** | **Tail**<br>**somites** | **Total**<br>**somites** |
| --- | --- | --- | --- |
| 1 | 47 | 83 | 130 |
| 2 | 48 | 84 | 132 |
| 3 | 46 | 89 | 135 |
| 4 | 49 | 90 | 139 |
| 5 | 47 | 84 | 131 |
| 6 | 46 | 85 | 131 |
| 7 | 46 | 88 | 134 |
| 8 | 46 | 98 | 144 |
| 9 | 48 | 90 | 138 |
| 10 | 48 | 86 | 134 |
| 11 | 49 | 84 | 133 |
| 12 | 46 | 98 | 144 |
| 13 | 48 | 89 | 137 |
| 14 | 50 | 89 | 139 |
| 15 | 48 | 90 | 138 |
| 16 | 52 | 86 | 138 |
| 17 | 49 | 89 | 138 |

**Table S2.** Counts of trunk and tail vertebrae in hatchling skates.

|  | **Trunk**<br>**vertebrae** | **Tail**<br>**vertebrae** | **Total**<br>**vertebrae** |
| --- | --- | --- | --- |
| 1 | 48 | 101+ | 150+ |
| 2 | 48 | 107+ | 152+ |
| 3 | 47 | 111+ | 158+ |
| 4 | 49 | 108+ | 157+ |
| 5 | 47 | 99+ | 146+ |
| 6 | 49 | 104+ | 153+ |
| 7 | 49 | 107+ | 156+ |
| 8 | 48 | 97+ | 145+ |

